# OpenGerminal: an open-source implementation of the Germinal antibody design pipeline

**DOI:** 10.64898/2026.06.25.734527

**Authors:** Bing Han, Sheng Li

**Affiliations:** Center for Membrane and Cell Physiology, University of Virginia, Charlottesville, VA, USA; Department of Molecular Physiology and Biological Physics, University of Virginia School of Medicine, Charlottesville, VA, USA; School of Data Science, University of Virginia, Charlottesville, VA, USA

## Abstract

Germinal is a recently described computational pipeline for de novo antibody design that combines AlphaFold-Multimer hallucination with antibody language model guidance to generate epitope-targeted antibodies. Germinal identified binders with nanomolar-to-low-micromolar affinities by testing only 43–101 designs per target across four diverse antigens, establishing it as a practical tool for epitope-directed antibody design accessible to standard academic laboratories. As this architecture is itself very recent, systematic replacement and benchmarking of its individual components remains largely unexplored, yet offers a valuable opportunity to probe the robustness of the underlying design. We present OpenGerminal, which replaces PyRosetta with a fully open-source stack comprising OpenMM 8.5.1, FreeSASA, FASPR, Biopython, and sc-rs v1.0.0, and adopts AbLang1 (ablang2 v0.2.1) as the sole antibody language model in place of IgLM. Benchmarking on two VHH targets (PD-L1 and IL-3) reveals that OpenGerminal achieves a markedly higher cofolding pass rate (PD-L1: 33.7% vs. 18.6%; IL-3: 24.6% vs. 8.0%) with equivalent or improved Chai-1 structural confidence metrics in accepted designs, at the cost of a modest increase in per-trajectory computation time (≥1.5×). Multi-chain target support is also extended and verified to run without error on the official insulin example. OpenGerminal provides the first systematic benchmarking of IgLM versus AbLang1 within the Germinal architecture, and its fully open-source component stack broadens the range of deployment contexts in which the pipeline can be used.

**Availability and Implementation:** Source code is freely available at https://github.com/teaninja/OpenGerminal under the Apache 2.0 license. Persistent archives with DOIs are available at https://doi.org/10.5281/zenodo.20755400 (code) and https://doi.org/10.5281/zenodo.20756013 (container image). OpenGerminal is implemented in Python and distributed as an Apptainer container (opengerminal_v1.0.0.sif). Installation instructions and example configurations are provided in the repository.

**Supplementary information:** Supplemental Figure 1 is available online.

## 1 Introduction

The computational design of antibodies and antibody fragments remains one of the most challenging problems in protein engineering. Traditional antibody discovery relies on hybridoma technology, phage display, or immunization campaigns, which are time-consuming, resource-intensive, and offer limited control over the target epitope. The emergence of deep learning-based protein design tools has begun to transform this landscape. Diffusion-based methods exemplified by RFdiffusion (Watson, et al., 2023) have demonstrated the ability to design high-affinity protein binders with exceptional efficiency, and RFdiffusion has more recently been extended to the antibody design problem, producing designs with atomic accuracy and experimental validation at nanomolar affinity (Bennett, et al., 2026). However, translating computational designs into experimentally confirmed binders typically requires high-throughput screening of large candidate pools, a logistical and infrastructural demand that places these approaches out of practical reach for many research groups. A complementary paradigm, hallucination-based design, backpropagates through structure-prediction networks such as AlphaFold2 to generate candidate binders, which—combined with structural quality filters— achieve high experimental success rates without large-scale screening; BindCraft (Pacesa, et al., 2025) exemplifies this approach. Germinal (Mille-Fragoso, et al., 2026) extends this hallucination paradigm to antibody design by combining gradient-based hallucination through AlphaFold-Multimer (Evans, et al., 2022) with sequence naturalness constraints from an antibody-specific language model, while anchoring designs to a fixed antibody framework to preserve antibody-like properties including robust mammalian cell expression. The original implementation employs IgLM (Shuai, et al., 2023) as the antibody language model (Mille-Fragoso, et al., 2026). Experimental testing of 43–101 designs per target across four antigens identified binders with nanomolar-to-low-micromolar affinities in all cases, with BLI-validated hit rates of 4–10% across all tested designs—comparable to substantially more complex pipelines (Swanson, et al., 2025). These results establish Germinal as a practical and effective tool for epitope-directed antibody design accessible to standard academic laboratories. Despite this promise, the broad utility of Germinal as originally published is constrained by two of its dependencies. PyRosetta (Chaudhury, et al., 2010) and IgLM are distributed under restrictive licenses—commercial and non-commercial, respectively—that limit unrestricted use and redistribution of the pipeline. Removing these constraints would broaden the contexts in which the method can be applied and reproduced. While AbLang1 (Olsen, et al., 2022) has since been integrated into the Germinal codebase as an alternative language model option (PR#55), the published Germinal pipeline reports results exclusively with IgLM and no comparative performance data between the two models have been published; notably, the authors themselves acknowledge IgLM’s restrictive license as a limitation and suggest open-source alternatives as a direction for future development (Mille-Fragoso, et al., 2026). Germinal is also a recently developed pipeline whose component interactions are not yet fully characterized; replacing and benchmarking individual modules provides an important opportunity to probe the robustness of the underlying architecture. Here we present OpenGerminal, an open-source implementation of the Germinal pipeline in which PyRosetta is replaced with an open-source computational stack and AbLang1 is adopted as the sole language model in place of IgLM.

## 2 Implementation

### 2.1 Overview of the Germinal pipeline

Germinal operates in four stages (Figure 1A). Stage 1 (sequence design) employs gradient-based hallucination through ColabDesign’s AlphaFold-Multimer interface (Frank, et al., 2024; Ovchinnikov, 2021), optimizing CDR sequences via PCGrad gradient merging of structural (pLDDT, iPTM, PAE) and sequence naturalness (antibody language model log-likelihood, lm_ll) objectives. Stage 2 (initial cofolding filter) evaluates each hallucinated design by independent structure prediction with Chai-1 v0.6.1 (Chai Discovery team, et al., 2024) followed by structural relaxation and interface scoring; designs failing geometric thresholds are discarded. Stage 3 (CDR redesign) applies AbMPNN to diversify CDR sequences while preserving the hallucinated backbone; this stage is independent of both the antibody language model and the structural scoring module. Stage 4 (final filter) re-folds AbMPNN-redesigned sequences with Chai-1 and applies a stricter set of structural and physicochemical thresholds. The original Germinal paper used AF3 for cofolding and filtering; OpenGerminal uses Chai-1 v0.6.1, which is included in the Germinal container and freely available for academic and commercial use without separate installation.

**Figure 1.**
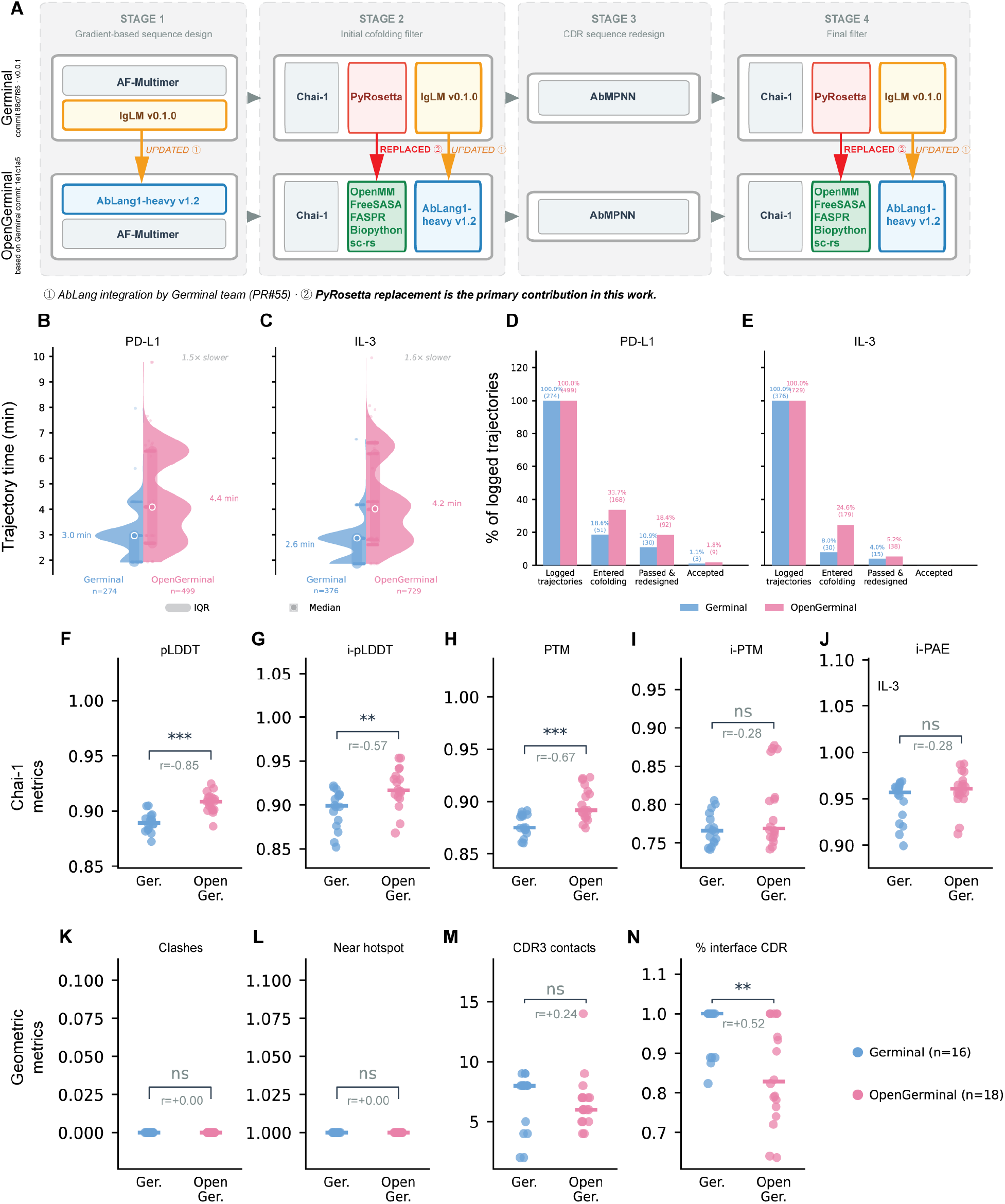
OpenGerminal pipeline architecture and benchmark performance against Germinal. **(A)** Schematic comparison of the Germinal (top row) and OpenGerminal (bottom row) pipelines across four stages. Stage 1: gradient-based sequence design using AF-Multimer hallucination guided by an antibody language model sequence loss. Stage 2: initial cofolding filter using Chai-1 structure prediction and interface scoring with structural relaxation. Stage 3: CDR sequence redesign with AbMPNN (unchanged in both versions). Stage 4: final filter applying Chai-1 confidence thresholds and interface geometry metrics. In OpenGerminal, IgLM v0.1.0 is replaced by AbLang1-heavy v1.2 (① integrated by the Germinal team via PR#55), and PyRosetta is replaced by an open-source stack comprising OpenMM, FreeSASA, FASPR, Biopython, and sc-rs (② primary contribution of this work). **(B–C)** Trajectory generation time (hallucination stage) for PD-L1 (B) and IL-3 (C) targets, shown as half-violin plots with individual data points. White-outlined circle indicates the median; shaded region indicates the IQR. Numerical labels indicate the group mean. Germinal: n = 274 trajectories for PD-L1, n = 376 for IL-3; OpenGerminal: n = 499 for PD-L1, n = 729 for IL-3. **(D–E)** Pipeline yield per logged trajectory for PD-L1 (D) and IL-3 (E). Bars show the percentage of trajectories reaching each stage: logged trajectories (denominator), entered initial cofolding filter, passed cofolding and entered CDR redesign, and final accepted designs. Accepted counts represent the number of unique design seeds, not the total number of accepted PDB structures (each seed may yield up to four AbMPNN sequence variants). All percentages are calculated relative to the number of logged trajectories (Germinal: n = 274 for PD-L1, n = 376 for IL-3; OpenGerminal: n = 499 for PD-L1, n = 729 for IL-3). **(F–J)** Chai-1 structure prediction confidence metrics for all accepted PD-L1 designs: overall pLDDT (F), interface pLDDT (G), PTM (H), iPTM (I), and interface PAE (J). Germinal: 16 PDB structures from 7 unique seeds; OpenGerminal: 18 PDB structures from 9 unique seeds. **(K–N)** Geometric interface quality metrics for all accepted PD-L1 designs computed with Biopython: number of steric clashes (K), binder proximity to target hotspot residues (L), CDR3–hotspot contact count (M), and fraction of interface residues located in CDRs (N). Germinal: 16 PDB structures from 7 unique seeds; OpenGerminal: 18 PDB structures from 9 unique seeds. Horizontal bars in strip plots indicate group medians. Statistical significance: ***P < 0.001, **P < 0.01, *P < 0.05, ns P ≥ 0.05 (two-sided Mann–Whitney U test); the value below each significance marker is the effect size (rank-biserial correlation r), where positive r indicates higher ranks for Germinal. PD-L1, programmed death-ligand 1; IL-3, interleukin-3; CDR, complementarity-determining region; IQR, interquartile range.

### 2.2 PyRosetta replacement

PyRosetta is invoked in Stages 2 and 4 (Figure 1A) for two distinct purposes: structural relaxation (FastRelax) and interface scoring (score_interface). OpenGerminal replaces both functions using an open-source stack, following the approach established in FreeBindCraft (Ring, 2025). Structural relaxation is performed with OpenMM 8.5.1 (Eastman, et al., 2017) combined with FASPR (Huang, et al., 2020) for side-chain repacking. Interface scoring employs FreeSASA (Mitternacht, 2016) for solvent-accessible surface area calculations, Biopython (Cock, et al., 2009) for geometric interface analysis, and sc-rs v1.0.0 (Ring, 2025) for shape complementarity following the Lawrence and Colman (1993) algorithm (Lawrence and Colman, 1993). All nine function interfaces of the original PyRosetta utility module are fully implemented, providing an API-compatible, PyRosetta-free backend with approximate replacements for the energy-derived metrics (see below). All modifications are incorporated into the distributed Apptainer container (opengerminal_v1.0.0.sif), which does not include PyRosetta.

Three interface metrics retain hard-coded placeholder values, as the corresponding PyRosetta- specific energy units cannot be directly mapped to OpenMM outputs without recalibration. The binder Rosetta energy score (binder_score, fixed at −1.0) and interface binding free energy (interface_dG, fixed at −10.0) are not applied as filter thresholds and are recorded in the output CSV for reference only. However, binder_score serves a second role in Stage 2: it is used by the ensemble selection logic to identify the best-relaxed structure from five candidates. Because the value is fixed, all five candidates receive identical scores and the first structure is always selected rather than the energetically optimal one; this represents a known degradation of the ensemble selection behavior, though its practical impact is expected to be limited given the typically small energetic differences between relaxed structures. The interface hydrogen bond count (interface_hbonds, fixed at 5) is applied as a filter threshold in Stage 4 (≥3 required); because the fixed value always satisfies this threshold, this filter is effectively disabled in OpenGerminal.

### 2.3 Antibody language model

The original Germinal pipeline uses IgLM as the antibody language model at three stages: in Stage 1 as a sequence naturalness loss guiding hallucination, and in Stages 2 and 4 to compute lm_ll, which is recorded in the output CSV but not applied as a filter threshold (Mille-Fragoso, et al., 2026). AbLang1 was subsequently integrated into the Germinal codebase as an alternative option (PR#55), but remained non-default and no comparative data were reported. OpenGerminal adopts AbLang1-heavy (ablang2 v0.2.1) as the sole language model, with IgLM removed as a selectable option. The language model substitution is not an original algorithmic contribution of this work; its effect on pipeline behavior is characterized empirically in Section 3.

### 2.4 Multi-chain target support

Germinal supports multi-chain target proteins as an experimental feature, but the original codebase contains several bugs that prevent the pipeline from completing when a multi-chain target is specified. OpenGerminal fixes three of these in modules shared by both pipelines: a KeyError triggered by comma-separated chain identifiers (filter_utils.py), a chain-renaming error in complexes with three or more chains (utils.py), and a sequence lookup failure for multi-chain targets during Chai-1 prediction (chai.py). A fourth fix to the PyRosetta interface string (pyrosetta_utils.py) is applied to the Germinal reference pipeline to enable benchmark comparison; this module is not invoked in OpenGerminal. All fixes are conditional on the presence of multiple target chains and do not affect single-chain runs. In the open-source scoring module (pr_alternative_utils.py), multi-chain support is further extended by correcting FreeSASA selection syntax and enabling shape complementarity scoring via temporary merging of target chains for sc-rs, which natively accepts only single-chain inputs.

## 3 Results

### 3.1 Benchmark design

To evaluate OpenGerminal against the original Germinal pipeline, we ran both versions on two previously reported targets: PD-L1 (programmed death-ligand 1) and IL-3 (interleukin-3), originally used to benchmark Germinal (Mille-Fragoso, et al., 2026). Both pipelines were run using the same target hotspot residues and VHH framework. Trajectory timing and pipeline yield were extracted from job logs; only trajectories with complete timing records were included (Germinal: n = 274 for PD-L1, n = 376 for IL-3; OpenGerminal: n = 499 for PD-L1, n = 729 for IL-3). Accepted design quality metrics were evaluated on all accepted designs identified from output structures (Germinal: 16 PDB structures from 7 unique seeds for PD-L1; OpenGerminal: 18 PDB structures from 9 unique seeds for PD-L1). IL-3 yielded no accepted designs in either pipeline.

### 3.2 Trajectory generation time

Measured at the hallucination stage (Stage 1; NVIDIA A100 80 GB GPU, UVA Rivanna/Afton HPC), OpenGerminal trajectories required a mean of 4.4 min for PD-L1 (vs. 3.0 min for Germinal, 1.5×) and 4.2 min for IL-3 (vs. 2.6 min, 1.6×) (Figure 1B–C). These figures represent a lower bound on the true end-to-end time difference: because OpenGerminal generates a substantially higher proportion of hallucinated sequences that pass the initial cofolding filter ( Section 3.3), a larger fraction of trajectories proceed to the computationally demanding Stages 2–4, further increasing total wall time per batch run.

### 3.3 Pipeline yield

A striking difference between the two versions emerged at the initial cofolding filter (Stage 2). OpenGerminal demonstrated markedly higher cofolding entry rates relative to Germinal: 33.7% vs. 18.6% for PD-L1 (1.81-fold increase) and 24.6% vs. 8.0% for IL-3 (3.08-fold increase) (Figure 1D–E). Cofolding pass rates followed the same trend (18.4% vs. 10.9% for PD-L1; 5.2% vs. 4.0% for IL-3). The proportion of accepted design seeds per logged trajectory was modestly higher for OpenGerminal for PD-L1 (1.8% vs. 1.1%), with both pipelines yielding zero accepted designs for IL-3. Because Stage 1 employs the same AlphaFold-Multimer model in both pipelines (verified by identical checksums of the core model code), and the antibody language model is the only other component active during hallucination, the observed differences in trajectory generation time and cofolding pass rates between Germinal and OpenGerminal are attributable to the substitution of IgLM with AbLang1.

### 3.4 Quality of accepted designs

We compared the structural quality of accepted PD-L1 designs between pipelines using Chai-1 confidence metrics and Biopython-derived geometric interface metrics (Figure 1F–N). Because both sets of metrics are computed using identical tools applied to structures from each respective pipeline, the comparison reflects genuine differences in design properties arising from the substitution of both the antibody language model (IgLM to AbLang1) and the structural relaxation protocol. OpenGerminal accepted designs showed significantly higher overall structure prediction confidence (pLDDT: median 0.908 vs. 0.889, p < 0.001, r = −0.85; PTM: 0.892 vs. 0.875, p < 0.001, r = −0.67) and interface prediction confidence (interface pLDDT: 0.917 vs. 0.899, p < 0.01, r = −0.57), while interface iPTM (0.769 vs. 0.766, r = −0.28) and interface PAE (0.961 vs. 0.957, r = −0.28) did not differ significantly (two-sided Mann–Whitney U test; effect sizes reported as rank-biserial correlation r, where negative values indicate higher ranks for OpenGerminal, in Figure 1F–N). All accepted designs from both pipelines were free of steric clashes, positioned near target hotspot residues, and showed similar CDR3–hotspot contact counts. The fraction of interface residues located in CDRs was modestly lower in OpenGerminal designs (median 82.8% vs. 100%, p < 0.01, r = 0.52), suggesting that the backbone hallucinations produced by the two pipelines favor slightly different interface geometries.

### 3.5 Post-hoc PyRosetta validation

To independently assess interface properties of OpenGerminal accepted designs using a common scoring framework, we applied PyRosetta scoring directly to the OpenMM-relaxed accepted structures without additional re-relaxation (Supplementary Figure 1A–I). Cross-validation of open-source pipeline metrics against PyRosetta scores confirmed high correspondence for shape complementarity (sc-rs vs. PyRosetta, Spearman ρ = 0.882) and good correspondence for interface hydrophobicity (FreeSASA vs. PyRosetta, ρ = 0.763) and surface hydrophobicity (ρ = 0.571), the latter showing a systematic upward scaling (slope = 1.20) that may reflect differences in surface definition between FreeSASA and PyRosetta. Backbone RMSD comparison (Biopython vs. PyRosetta) yielded a Spearman ρ of 0.591; Pearson correlation was near zero due to a single outlier design.

Application of the five PyRosetta-based Stage 4 filter thresholds to post-hoc scores showed that the distributions of all five metrics overlapped broadly between Germinal and OpenGerminal accepted designs, confirming that OpenGerminal designs fall within the same general quality range as Germinal designs. These comparisons should be interpreted cautiously: Germinal designs were scored after PyRosetta FastRelax as part of the pipeline, whereas OpenGerminal designs were scored on OpenMM-relaxed structures, introducing a systematic difference in starting conformation.

### 3.6 Multi-chain pipeline verification

To verify multi-chain target support, we ran OpenGerminal on insulin, the official multi-chain example distributed with Germinal (target chains A and B; binder chain C; hotspots A2, A21, B12, B16). Of 300 hallucination trajectories, 85 (28.3%) passed the initial cofolding filter, and four redesign candidates were produced from one trajectory seed. No accepted designs were obtained; all four candidates failed the binder_near_hotspot filter and exhibited low Chai-1 interface confidence (external_iptm: 0.35–0.42, threshold >0.74), indicating that the hallucinated binder did not converge on the target hotspot region. The original Germinal codebase contains chain-handling bugs that prevent pipeline completion on this input; OpenGerminal’s fixes are required for the pipeline to run. These results confirm that the multi-chain bug fixes are functional; achieving accepted designs on multi-chain targets remains a challenge for future work.

## 4 Conclusion

We have presented OpenGerminal, a fully open-source implementation of the Germinal antibody design pipeline in which PyRosetta is replaced by an open-source stack and AbLang1 is adopted as the sole antibody language model in place of IgLM. OpenGerminal preserves the core design logic of the original pipeline while enabling unrestricted use in academic and commercial settings.

Benchmarking on two targets provides the first comparative data for IgLM versus AbLang1 within this architecture: AbLang1-guided hallucination substantially increases the fraction of sequences that pass initial Chai-1 cofolding validation, likely reflecting differences in the sequence space constraints imposed by the two models. Accepted designs show equivalent or improved Chai-1 structural confidence metrics relative to Germinal designs.

Several limitations of the current implementation warrant attention. First, OpenGerminal is slower than the original pipeline at the hallucination stage (≥1.5–1.6× per trajectory), with additional time incurred by the higher cofolding entry rate. Second, three interface metrics retain hard-coded placeholder values, of which only interface_hbonds directly affects filter behavior; recalibration to OpenMM energy units is required for full equivalence. Third, the Stage 2 ensemble selection logic is degraded due to the fixed binder_score, though practical impact is expected to be small. Fourth, while OpenGerminal incorporates bug fixes that extend its applicability to multi-chain target proteins, comprehensive validation of this functionality remains to be completed; multi-chain support is also experimental in the original Germinal pipeline.

Future development will focus on replacing hard-coded fixed values with true OpenMM-derived energy estimates and recalibrating filter thresholds, improving the Stage 2 ensemble selection using an accessible structural metric such as interface delta-SASA, and extending validated multi-chain target support. Prospective experimental validation of OpenGerminal-designed binders will be needed to fully assess the impact of component substitution on design success rates. We anticipate that OpenGerminal will lower barriers to adoption of the Germinal framework and facilitate systematic exploration of alternative computational components within the antibody design pipeline.

## Acknowledgements

The authors thank the Germinal development team (Santiago Mille-Fragoso and colleagues) for open-sourcing their pipeline, the developers of FreeBindCraft (https://github.com/cytokineking/FreeBindCraft) for open-sourcing their PyRosetta-free binder design pipeline which informed the development of OpenGerminal, Dr. Anne K. Kenworthy (Center for Membrane and Cell Physiology and Department of Molecular Physiology and Biological Physics, University of Virginia School of Medicine) for comments on the manuscript. Large language model tools (Claude, Anthropic) were used to assist with code development; the authors take full responsibility for the correctness of all code. The authors acknowledge Research Computing at The University of Virginia for providing computational resources and technical support that have contributed to the results reported within this publication (https://rc.virginia.edu). The content is solely the responsibility of the authors and does not necessarily represent the official views of the National Institutes of Health.

## Author Contributions

Bing Han (Conceptualization [lead], Data curation [lead], Formal analysis [lead], Investigation [lead], Methodology [lead], Validation [lead], Visualization [lead], Writing—original draft [lead], Writing—review & editing [lead]); Sheng Li (Formal analysis [supporting], Methodology [supporting], Writing—review & editing [supporting]).

