## Supplementary Figure 1 for "OpenGerminal: an open-source implementation of the Germinal antibody design pipeline"

**FigureS1**

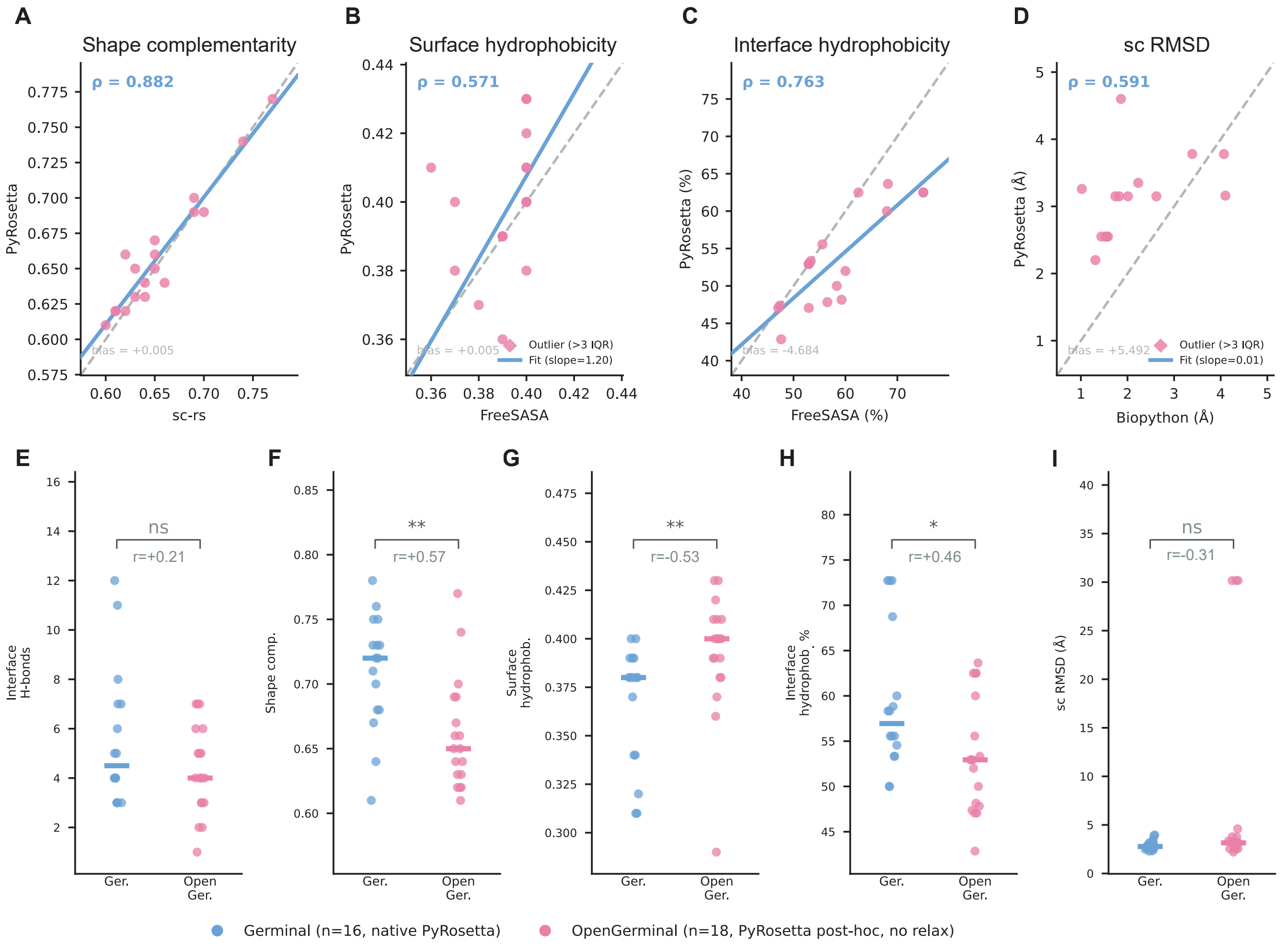
A
Shape complementarity
ρ = 0.882
bias = +0.005
PyRosetta
sc-rs
B
Surface hydrophobicity
ρ = 0.571
bias = +0.005
Outlier (>3 IQR)
Fit (slope=1.20)
PyRosetta
FreeSASA
C
Interface hydrophobicity
ρ = 0.763
bias = -4.684
PyRosetta (%)
FreeSASA (%)
D
sc RMSD
ρ = 0.591
bias = +5.492
Outlier (>3 IQR)
Fit (slope=0.01)
PyRosetta (Å)
Biopython (Å)
E
Interface H-bonds
ns
r=+0.21
F
Shape comp.
**
r=+0.57
G
Surface hydrophob.
**
r=-0.53
H
Interface hydrophob. %
*
r=+0.46
I
sc RMSD (Å)
ns
r=-0.31
Ger.
Open Ger.
Germinal (n=16, native PyRosetta)
OpenGerminal (n=18, PyRosetta post-hoc, no relax)

**Supplementary Figure 1. Post-hoc PyRosetta validation of OpenGerminal accepted designs for PD-L1. (A–D)** Cross-validation of open-source pipeline metrics against PyRosetta post-hoc scoring for OpenGerminal accepted designs (n = 18 PDB structures). Each point represents one accepted design. The dashed grey line indicates the identity line (y = x); the solid blue line shows the linear regression fit. Spearman rank correlation coefficient (ρ) is shown for each comparison. (A) Shape complementarity: sc-rs (pipeline) vs. PyRosetta (ρ = 0.882, slope = 0.90, bias = +0.005). (B) Surface hydrophobicity: FreeSASA (pipeline) vs. PyRosetta (ρ = 0.571, slope = 1.20, bias = +0.005). (C) Interface hydrophobicity: FreeSASA (pipeline) vs. PyRosetta (ρ = 0.763, slope = 0.62, bias = −4.684). (D) Structural RMSD between Chai-1 predicted structure and AF-Multimer hallucination backbone (sc RMSD): Biopython (pipeline) vs. PyRosetta (ρ = 0.591, slope = 0.01, bias = +5.492; one outlier exceeds the axis range and is not shown). PyRosetta scoring was performed directly on OpenMM-relaxed accepted structures without additional re-relaxation. **(E–I)** Comparison of PyRosetta interface metrics between Germinal accepted designs scored natively during the pipeline (n = 16 PDB structures from 7 unique seeds) and OpenGerminal accepted designs scored post-hoc without re-relaxation (n = 18 PDB structures from 9 unique seeds): interface H-bonds (E), shape complementarity (F), surface hydrophobicity (G), interface hydrophobicity (%) (H), and sc RMSD (I). Horizontal bars indicate group medians. Statistical significance: $^{**}P < 0.01$, $^{*}P < 0.05$, ns $P \geq 0.05$ (two-sided Mann–Whitney U test); the value below each significance marker is the effect size (rank-biserial correlation r), where positive r indicates higher ranks for Germinal. sc RMSD, structural RMSD between Chai-1 prediction and AF-Multimer hallucination backbone.
